# Bioschemas Training Profiles: A set of specifications for standardizing training information to facilitate the discovery of training programs and resources

**DOI:** 10.1101/2022.11.24.516513

**Authors:** Leyla Jael Castro, Patricia M. Palagi, Niall Beard, Teresa K. Attwood, Michelle D. Brazas

**Affiliations:** ZB MED Information Centre for Life Sciences; SIB Swiss Institute of Bioinformatics, Switzerland; Department of Computer Science, The University of Manchester, UK; Ontario Institute for Cancer Research, Toronto, CA

## Abstract

Stand-alone life science training events and e-learning solutions are amongst the most sought-after modes of training because they address both point-of-need learning and the limited timeframes available for ‘upskilling’. Yet, finding relevant life sciences training courses and materials is challenging because such resources are not marked up for Internet searches in a consistent way. This absence of mark-up standards to facilitate discovery, re-use and aggregation of training resources limits their usefulness and knowledge-translation potential. Through a joint effort between the Global Organisation for Bioinformatics Learning, Education and Training (GOBLET), the Bioschemas Training community and the ELIXIR FAIR Training Focus Group, a set of Bioschemas Training profiles has been developed, published and implemented for life sciences training courses and materials. Here, we describe our development approach and methods, which were based on the Bioschemas model, and present the results for the three Bioschemas Training profiles: *TrainingMaterial, Course* and *CourseInstance*. Several implementation challenges were encountered, which we discuss alongside potential solutions. Over time, continued implementation of these Bioschemas Training profiles by training providers will obviate the barriers to skill development, facilitating both the discovery of relevant training events to meet individuals’ learning needs, and the discovery and re-use of training and instructional materials.

## Introduction

Research outputs come in numerous formats, including scholarly publications, software, data, workflows, and training courses and materials. Although scholarly publications have long been recognized as the main and official output of research activities, the value of other research outputs such as data, workflows and software are now gaining recognition. Accordingly, efforts to make these other scientific outputs ‘Findable, Accessible, Interoperable and Reusable (FAIR)’^1,2–3,4^ and to improve ‘Research Data Management’^5^ have increased in recent years. As a result, new repositories and registries have emerged: for data (e.g., [Dryad](https://datadryad.org/), RE3Data), for research software (e.g., [Software Heritage Foundation archive](https://www.softwareheritage.org/)) and for workflows (e.g., [Dockstore](https://dockstore.org/), [WorkflowHub](https://workflowhub.eu/)).

Developing similar repositories or registries for training courses and materials has proven more challenging. Specialized e-learning platforms like [FutureLearn](https://www.futurelearn.com/), [Coursera](https://www.coursera.org/) exist, but are not inclusive of training activities outside their membership organizations. Similarly, institutional portals like [EMBL-EBI Training](https://www.ebi.ac.uk/training/) host training resources developed by institutional members. Wider efforts to collate training resources or announcements of training events, such as GOBLET’s Training Portal^6^, the iAnn.pro project^7^ or ELIXIR’s Training e-Support System TeSS^8^, have not taken hold, primarily because training providers need to register or upload their resources, a step too time-consuming for many to complete. In TeSS, for example, development of custom scripts to ‘scrape’ training information from different providers’ websites, and creation and maintenance of bespoke API clients is required. As a result, it is more difficult for people and search engines to find, aggregate, build upon and re-use training resources.

Yet in the life sciences, there is a growing need for bioinformatics, data science and computational training. Stand-alone events and e-learning solutions are amongst the most sought-after modes of training delivery, largely owing to the limited time that most individuals have available for ‘upskilling’, and because most training is sought at the point of need^9,10^. A standardized way to mark-up and thus discover, collate and analyze these training courses, offerings and training materials is badly needed.

In this paper, we present results from a joint effort between GOBLET^11^, the Bioschemas Training community^12^ and the ELIXIR FAIR Training Focus Group^13^ to devise, and facilitate implementation of, structured (meta)data for training courses and materials. We broadly introduce efforts to create standards, then focus on our approach and methods based on the Bioschemas working group model, and present the results for three Bioschemas Training profiles: TrainingMaterial, Course and CourseInstance. We conclude with a discussion around the implementation challenges and possible ways to move forward.

## Methods: Approaches to Standards Development for Training

### Schema.org

Schema.org^14^ provides a set of vocabularies to describe a vast array of entities on the Internet (films, events, people, organizations, *etc*.). The vocabularies comprise properties that define specific entity attributes: *e.g*., for a film, who the director is; for a course, when the start date is; for a person, their place and date of birth. These properties and their attributes, can be used by developers to annotate their websites, allowing search engines to better understand their content, and more effectively connect searchers with information sequestered in their pages. For example, Schema.org-annotated content may be promoted in ranked search results, or used to generate ‘knowledge panels’ (like those typically provided by Google searches). Vocabularies can be implemented on websites using three formats: RDFa, Microdata or JSON-LD, the latter being the most popular.

Although initially established by a consortium of search engines^14^, the Schema.org vocabularies are, in practice, finessed via community consensus. The process for developing extensions and incorporating them into the main vocabulary is well-documented, thereby facilitating contributions from a community.

At the heart of our project was the need to collate training-related data for widely dispersed communities of life-scientists. Specifically, we needed a schema that would facilitate discovery of marked-up resources (courses, workshops, training materials), that could be readily implemented with few technical barriers in a structured data format, making the data easy for aggregators to extract, load and transform. It was expedient to build on an existing standard rather than to reinvent one, as producing competing standards hinders interoperability.

As a starting point, we began with the Learning Resource Metadata Initiative (LRMI)^15^, which provides a set of schema definitions for marking up and describing educational resources, built on the Schema.org vocabularies and other standards. The LRMI work embodied in Schema.org afforded the best fit for our purposes: it has a simple mechanism for implementation, provides properties well-aligned with our needs, and has a tool ecosystem that makes extraction and re-use of annotated data straightforward. However, as Schema.org vocabularies are Internet-wide, they attempt to fulfill a wide variety of use-cases - practically every industry uses them. As such, their properties are often bloated with niche terms that are not relevant to the life sciences. Other efforts, such as those from the Australian Research Data Commons^16^ and Research Data Alliance^17^, were also used, although neither has described an implementation path.

To leverage the benefits of Schema.org and these other initiatives, and to address their limitations for the bioinformatics training community, the Bioschemas Training profiles initiative emerged to customize and propose amendments to Schema.org vocabularies, specifically to encompass training aspects in the life sciences. Here, we present the status of the Bioschemas Training initiative, and invite further community input and adoption.

### Bioschemas development approach

The Bioschemas approach to create schema specifications puts life science research communities at the center. A research community in a specific subject or field (*e.g*., proteins, chemicals, training) leads the schema development. Whenever a community identifies the need to have common metadata to describe elements or activities relevant to them, this community can use the Bioschemas approach to define, share, validate and publish a schema specification. Based on Schema.org, Bioschemas defines two possible schema specifications: types and profiles. A Bioschemas type corresponds to something that does not have a type in Schema.org, such as *Protein* or *Gene*, both of which were types proposed by the Bioschemas community and have been accepted into the pending section of Schema.org. A Bioschemas profile corresponds to a usage recommendation for an existing Schema.org type, such as Bioschemas profile for Schema.org type *Dataset* or *TrainingMaterial*. In Schema.org, the type *Dataset* has more than 100 properties (*i.e*., relations to other types that help describe a *Dataset*, such as its abstract or its authors), making it difficult for researchers to figure out which properties are the most relevant to use. To overcome this limitation, Bioschemas profiles are proposed and supported by communities.

The Bioschemas development approach to create profiles is provided as a tutorial on the Bioschemas website^18^, and can be summarized as follows:

- A community identifies an element or activity that requires a common set of metadata. The community group defines the overall objectives, and names its leaders and members. This group is charged with the creation (and subsequent maintenance) of the profile.
- The community group defines the use cases for the profile (*e.g*., information exchange, aggregation, summarization).
- The group conducts a search, identifies relevant metadata schemas (even if not formalized) and creates a ‘crosswalk’. A crosswalk is a compilation of the existing metadata schemas identifying properties, synonyms, similarities and differences; it also includes the properties relevant to the community. The idea behind a crosswalk is to get a better understanding of the community needs, current offerings and gaps.
- Based on the community needs and use cases, the group defines the marginality assignments for each property according to community usage. Properties are assigned minimum (*e.g*., mandatory), recommended (*e.g*., not always available, but when they are, they should be provided) or optional (*e.g*., nice to have) marginality. The group also defines the cardinality - one or many - for each property (*e.g*., a *CourseInstance* has one description only, but can have many instructors).
- The group searches for a compatible type in Schema.org. A compatible type is one that already corresponds to the element or activity to be described. For instance, Schema.org *Course* is the type used for the Bioschemas *Course* profile.
- If no Schema.org type is identified, the group proposes a new type. The approach to create and propose new types is out of the scope of this paper, as training profiles already existed in Schema.org.
- If a compatible Schema.org type is identified, the group uses a spreadsheet, including all the properties for this Schema.org type. Bioschemas provides a template for such spreadsheets.
- The community group selects those properties corresponding to the identified needs (from the crosswalk) and either adopts the use as is from Schema.org, or adds a Bioschemas description, if further clarification on how to use this property is necessary.
- The group identifies those properties where controlled vocabularies can be used (*e.g*., for disambiguation purposes). A Controlled Vocabulary (CV) can be a recommended list of free-text terms (*e.g*., ‘beginner’, ‘intermediate, ‘advanced’), an enumeration (*e.g*., https://schema.org/EventAttendanceModeEnumeration, which includes mixed, offline and online attendance modes), a standard (*e.g*., currency codes, such as USD, EUR or GBP), or an ontology (*e.g*., EDAM^20^).
- The group characterizes the properties with respect to marginality (*e.g*., minimum, recommended or optional) and cardinality (one, many) as per the crosswalk exercise.
- At this point, the profile is ready for publication. The spreadsheet is published to the Web (an option provided by Google spreadsheets) and converted to YAML format^19^ (via the command interface application provided by Bioschemas). The YAML format is integrated into the Bioschemas website by creating a Pull Request for the new profile.
- All new profiles (or updates of an existing one) are considered draft. For a profile to become a released profile, the group calls for a community consultation, where anyone can participate in a discussion to improve or accept the profile as proposed. A minimum of two adopters (*i.e*., two independent organizations using the profile to mark-up their Web pages) is also required.

### The training community development of Bioschemas Training profiles

A group of training providers, training networks and related organizations (initially encompassing ELIXIR, GOBLET, Pistoia Alliance, and later joined by EMBL-EBI, SIB Swiss Institute of Bioinformatics, Bioinformatics.ca, the Carpentries, etc.) came together in 2015 as a community to establish standards for training resources. When the Bioschemas initiative launched, this training community formed the Bioschemas community tasked with defining the Training Profiles in Bioschemas.

The Schema.org vocabularies most suited to helping define training were identified, and their properties assessed. The group also reviewed the metadata exposed on each of their own websites, and compared their properties against those of the relevant Schema.org vocabulary. To reduce the complexity of the Schema.org vocabularies, we focused on the applicable training-specific vocabulary and created profiles for them. The motivation for creating Bioschemas Training profiles was to supplement the existing Schema.org specifications, thereby improving their utility for life scientists (*e.g*., increasing the interoperability of resources annotated with the profiles, and improving their usability, by offering more support, examples and guidance for adopters).

Following the Bioschemas development approach above, our community group determined for each property:

1. Whether the property was used across the metadata schemas of our training community websites. This involved identifying properties that appeared to be equivalent, despite using different terminology (*e.g*., the *keyword* property could be referred to as *tags* or *keywords* on different provider websites, or the *contributor* property could be referred to as *contributor, submitter* or *maintainer);*
2. The prevalence of the property’s use across the different websites to guide marginality assignments (*e.g*., the property *name* was used in all the websites, while the *contributor* property was used in 70% of them). To help determine marginality categorisations, LRMI authors^15^ were consulted;
3. The expected property value *types (e.g*., an *author* could be an *Organization* or a *Person;* a *version* could be a *Number* or *Text;* an *identifier* could be a *URL* or *Text*);
4. Whether the property values could be constrained by an existing ontology or standard (*e.g*., a topic term could be mapped to the EDAM ontology^20^, or a course duration could be formatted with reference to the ISO8601 standard^21^), or in the absence of an existing ontology or standard, whether a custom vocabulary would be helpful;
5. Whether there were properties used in provider websites that weren’t included in the Schema.org vocabularies; and
6. Whether the property’s description required more specific instructions or interpretations for usage by a life scientist.

The resulting matrix of properties formed the basis of three Bioschemas training profiles: *Course*, *CourseInstance* and *TrainingMaterial*.

To further refine the draft Training profiles, the Training community group held several in-depth reviews at the 2017 GOBLET AGM in Lisbon, the 2018 GOBLET AGM in Toronto, at Biohackathon Europe 2018 in Paris, and at the 2019 Bioinformatics Education Summit in Cape Town. A series of follow-up discussions were also held online. During these reviews, participants were given a list of considerations with which to ensure that the standard being produced conformed with, and supported, the use-cases. For each profile, thought was given to whether: a) its properties should be regarded as minimum (essential), recommended, optional or irrelevant for our purposes; b) its property values could be constrained by a CV; and c) an additional description was necessary to guide the user. As per the Bioschemas process, our decision-making process was guided by the following Bioschemas definitions:

*Minimum properties*

1. are essential for helping searchers to discover a training resource(s);
2. must be satisfied if resources are to be Bioschemas compliant (if a minimum property cannot be satisfied by a provider, then the resource doesn’t comply with the standard);
3. should be small in number, and ubiquitous across providers. There should be an absolute maximum of 6 minimum properties.

*Recommended properties*

1. are not essential, but nevertheless provide additional information to facilitate discovery of resources, and help searchers make decisions about their relevance;
2. have high priority relative to optional properties.

*Optional properties*

1. provide additional information that might enhance the granularity of a search;
2. are unlikely either to be available from all providers, or to be used by searchers to find resources, but might help them to determine their relevance;
3. have relatively low priority.

*Unnecessary properties*

1. are unlikely either to be employed by providers, or to help searchers to find resources or make decisions about their relevance.

*Controlled Vocabularies*

1. are beneficial in making heterogeneous content comparable;
2. are not equally well-established or widespread, so should be recommended advisedly;
3. if custom-made, must cover every potential value that a provider may require, be maintained over time, and be mapped to other CVs, if they become available.

*Descriptions*

1. are understandable and applicable to life science users;
2. if new descriptions are added, they are tailored to life science users.

This iterative review process yielded a set of draft specifications for each Training profile. These were submitted to the Bioschemas governance review panel and remained as drafts until each profile achieved two adopters, after which the profiles were released into current specifications.

## Results

The series of international community reviews led to the development of three Bioschemas Training profiles, each of which is interlinked (Figure 1). For describing courses, new Bioschemas profiles were created for *Course* and *CourseInstance* (derived from the *Schema.org/Course* and *Schema.org/CourseInstance* vocabularies, respectively, as defined by the LRMI). For describing training materials, a new Bioschemas profile was created for *TrainingMaterial*, derived from *Schema.org/CreativeWork/LearningResource* vocabulary.

**Figure 1.**
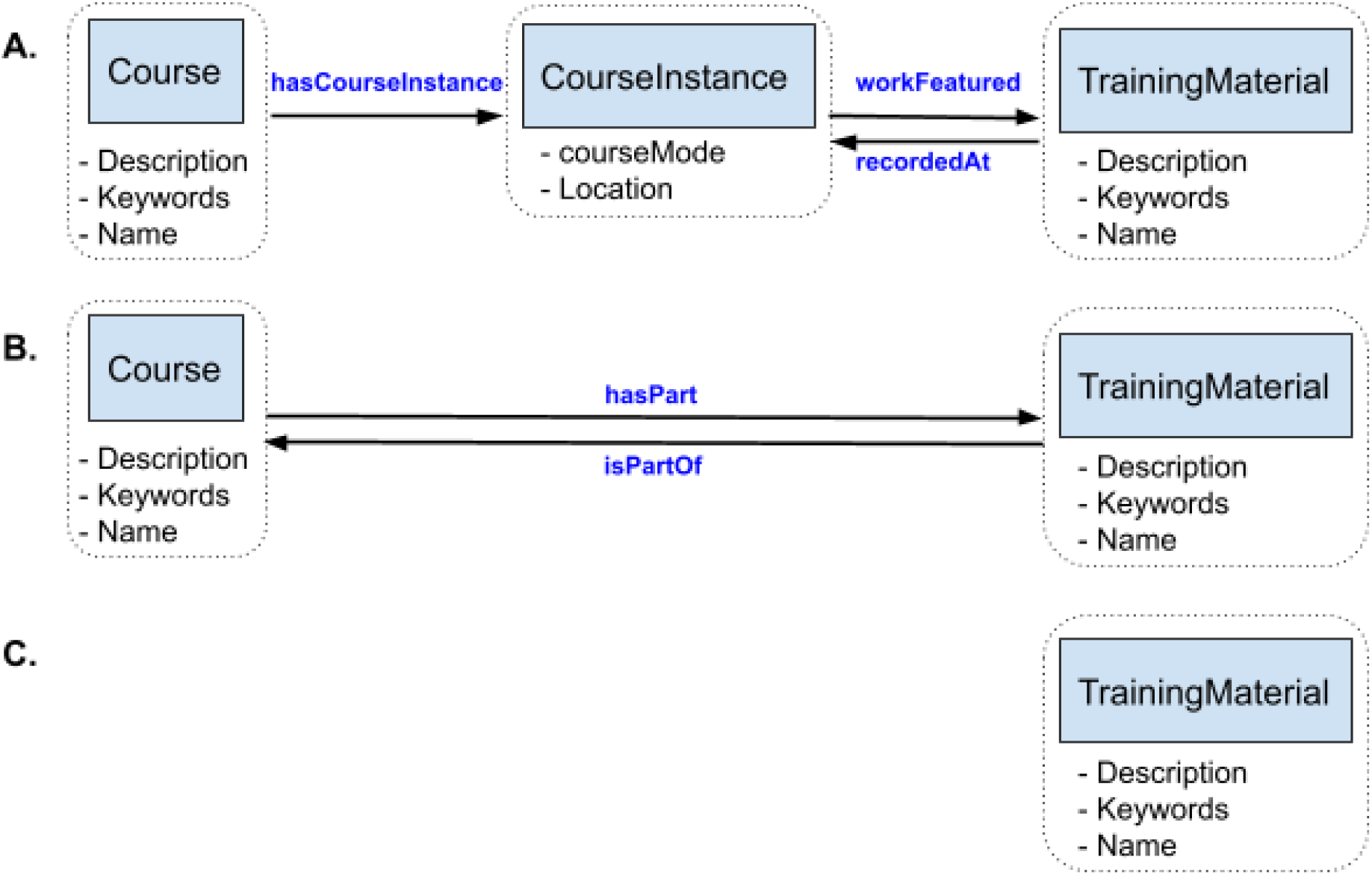
Diagram showing how the Training profiles for *Course, CourseInstance* and *TrainingMaterial* interconnect with each other through various properties (blue text). The minimum properties ascribed to each profile are listed within the dashed boxes. A) describes how to link *TrainingMaterial* within a *CourseInstance* for a *Course*, such as for multiple calendar offerings of a course. B) describes how *TrainingMaterial* can be associated with a *Course* that does not have a *CourseInstance*, such as for a self-directed online course. C) describes stand-alone *TrainingMaterial*, such as a recorded presentation.

### The specifications and their linkage

*Course* represents the concept of any training course, workshop, conference event or e-learning resource. It defines the content and attributes of the course (including its name, description, learning outcomes, duration, *etc*.), information about who is providing it (*e.g*., provider, authors, contributors), and the intended audience. For example, an organization may have developed a half-day *Course* for teaching R for a general audience, and named it “Introduction to R”.

*CourseInstance* represents a specific course offering scheduled at a given time or place. Multiple *CourseInstances* can be described for any given *Course*. The *Course-CourseInstance* paradigm allows training providers to describe repeated offerings of the same course without duplicating other metadata. This is useful for short courses that are offered periodically, or that run in different locations. Several properties defined in *Course (e.g., duration, description, name*) can be overridden in *CourseInstance* to allow for variations in a given course offering. In the above example, the general *Course* for teaching R is agnostic of the audience (*e.g*., “Introduction to R”). If this generic *Course* is customized for geoscientists, for example, the *name* property within the root R *Course* may be overridden with the *CourseInstance name* property to indicate a geoscience-focused course offering (*e.g*., “Introduction to R for Geoscientists”). To ensure that a *CourseInstance* is always associated with a *Course*, the *Course teaches* property, which describes the learning outcomes, is not a property of *CourseInstance*.

*TrainingMaterial* represents learning resources such as books, videos, lectures, tutorials, slide-decks, articles, and so on, which can be linked (where relevant) to their cognate training events, as per Figure 1. Continuing with the above example, a reference sheet of common R functions is developed and this training material is used by both the “Introduction to R” and “Introduction to R for Geoscientists” courses.

The specifications are designed to be used in conjunction with one another, allowing their relationships and interconnections to be expressed explicitly, as illustrated in Figure 1. Thus, for example, a *Course* may point to a variety of *CourseInstances*, and to its associated *TrainingMaterials* (slides, hand-outs, *etc*.), and *TrainingMaterials* may come from either a *Course* or a *CourseInstance*, and so on. Relevant profile relationships to point out include:

- *Course* can be linked to *CourseInstance* through the *has CourseInstance* property, and to a *TrainingMaterial* through the *hasPart* property;
- *TrainingMaterial* can link to the *Course* to which it belongs via the *isPartOf* property or to a *CourseInstance* of which it is a part through the *recordedAt* property.

### The profiles and their minimum properties

The full set of specifications is represented in tabular format, grouped according to the marginality categories: minimum, recommended and optional. The first section sets out the minimum properties, the number of which is kept as small as possible to encourage more adopters and avoid becoming a barrier to adoption.

*Course* and *TrainingMaterial* have the same minimum properties: *description, keywords* and *name*. These are present in the existing Schema.org specifications, where their descriptions were deemed suitable for the life science training community. The cardinality (whether a property can occur only once or multiple times) is set to *one* for each property. This means that implementations of the schema must have exactly one of each of these properties: one name, one description, one set of keywords. Note that the latter property - *keywords* - is pluralized to allow for single instances of comma-separated lists containing multiple keywords.

*CourseInstance* has minimum properties *courseMode* and *location*. The *courseMode* property can be of type either *Text* or *URL*. *courseMode* can be defined by several terms describing: i) the mode of training delivery (online, onsite, hybrid or blended); ii) the mode of training interaction (synchronous or asynchronous); and iii) the mode of study (full-time or part-time), chosen from vocabularies defined by the Common Education Data Standard^22^ or the Glossary of terms defined by the Bioschemas Training community^23^. The *location* property can be of type *Place, PostalAddress, Text or VirtualLocation*.

The complete set of specifications can be viewed on the Bioschemas website (https://bioschemas.org/groups/Training). Included are examples of usage for each profile and its properties, as well as the lastest deployments of the specifications, which serve as illustrations to those who wish to implement the profiles for their own courses and training materials. Supplementary Data contains figures illustrating how implementation of the profiles improves the machine readability of web posted training resources.

### A practical example of how to deploy a Training profile

From a course provider perspective, the only step that is required to markup webpages with these profiles is to include the profile descriptors either in JSON-LD, Microdata or RDFa format in the HTML code of the webpage.

Figure 2 gives an example of the JSON-LD format markup for the Course profile for a course entitled “Single-Cell Transcriptomics”. Here, only the minimal descriptors *name, keywords* and *description* are shown and contain the following:

- *name* – Single-Cell Transcriptomics
- *keywords* – next generation sequencing, single-cell biology, transcriptomics, DNA, RNA, epigenome, differential analysis, gene expression profiling
- *description* – This 3-day course covers the main technologies and aspects to consider while designing a scRNAseq experiment. In addition, it will cover the theoretical background of analysis methods with hands-on practical data analysis sessions applied to droplet-based methods.

**Figure 2.**
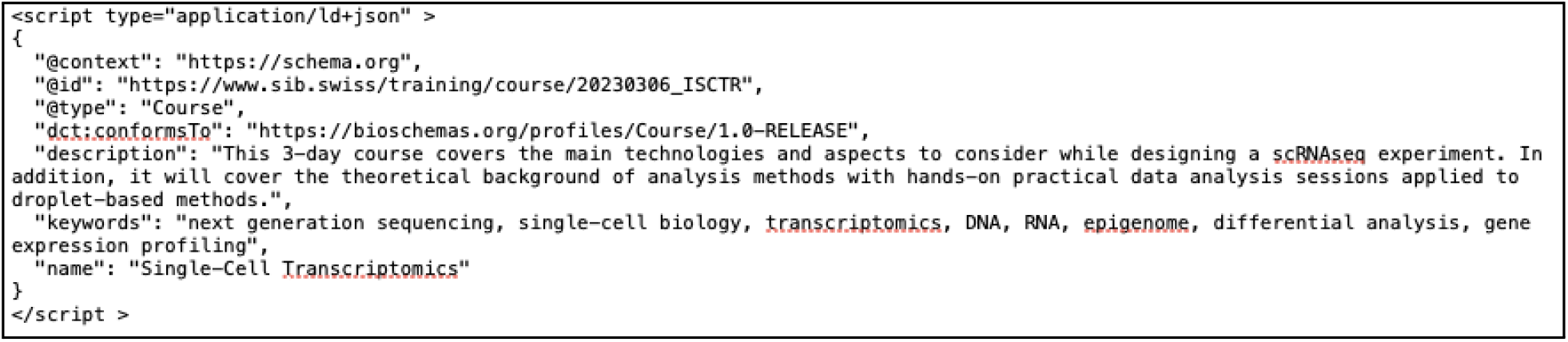
JSON-LD illustration for a Course entity, displaying standard and minimum properties.

Four other standard and mandatory properties makeup the JSON-LD markup:

- @context – states the use of schema
- @type – identifies the type of item being described, in this case, a Course
- @id – the identifier of the item being described, in this case the URL of the bespoken course
- dct:conformsTo – the Bioschemas profile version of the item

At a minimum, implementing a Bioschemas Training profile requires the set of properties listed under minimum; other properties can be incorporated where appropriate to the user. Implementation can then be validated using a variety of validation tools. The Bioschemas website provides a profile generator (https://github.com/BioSchemas/BioschemasMarkupGenerator) where this and any of the Bioschemas profiles can be easily generated before being inserted into a webpage.

## Discussion

### Drivers for and barriers to standards development and implementation

The Bioschemas community group for Training profiles set out to create schemas that could be implemented in a structured format. The goal was to provide specifications that would not be technically difficult to implement, that would facilitate discovery of appropriately marked-up life-science training resources, and would consequently be easy for aggregators to extract, load and transform.

The success of a standard depends on a variety of factors, but two phases are vitally important: development and proliferation. If a standard already exists, a new one doesn’t need to be produced, unless the existing standard fails to match the use-case(s) of interest, and its developers are unable, or unwilling, to participate in their further enhancement. Ultimately, standard development needs to be driven by real use-cases for which the standard seeks to provide solutions. To formulate helpful use-cases, it is necessary to include perspectives from a wide range of potential adopters, consensus from whom is necessary to validate the emerging standard. This can be difficult to achieve, and disagreements can stall the process if a procedure for resolution isn’t defined, and convincingly and consistently applied.

Once a standard begins to emerge, its proliferation depends on the burden of its technical implementation measured against the benefits perceived by potential adopters. As such, adoption by target communities depends on making those benefits clear, and lowering the conceptual and technical barriers sufficiently to make the adoption process as simple as possible. Wide participation in the standard-development phase, particularly buy-in from established and respected stakeholders, helps to expand the network available for disseminating the standard and for supporting its adoption. From the outset, we therefore engaged and sought input from a spectrum of training providers from across the globe.

An important conceptual hurdle arose during this process. The empirical assessment by the reviewers was based on how training courses and materials are currently described or used. Thus the marginalities they determined were indicative of how training information has been represented up to now. However, this may not be optimal future practice. We were therefore faced with the tension between developing a standard reflective of what training providers do now versus developing a standard that would advance best practices in the future, but importantly, without setting the barriers to adoption too high. For example, statements of learning outcomes, and citations of others’ work, were not routinely discovered across the websites under review; both, however, are clearly desirable. It was reasoned that setting their marginalities to Optional would encourage current practice, allowing such important features to continue to be overlooked. Conversely, setting them to Minimum might discourage adoption, as this would require more work from providers. By way of compromise, their marginalities were therefore denoted Recommended.

A second hurdle encountered during implementation was the requirement to upgrade the logic and backend of existing, often complex websites to introduce the Bioschemas Training profiles on large amounts of historical content, while also ensuring future content has the correct minimum metadata. For many providers, mapping their existing fields to the Bioschemas properties proved difficult, especially if a minimum property was missing within their websites. The implementation of the schema, however, is current and forward looking, such that minimum properties should be populated for all current and future courses and materials. For other providers, who outsource their website development to external companies and are charged for feature implementation, updating their websites with a new schema is not a high priority, given the financial cost to do so, despite the many advantages of having a schema. As a result, adoption of each Training profile has taken many years, far longer than anticipated when the community embarked on their development.

### Advantages and Challenges of the Bioschemas specifications

The Schema.org vocabularies are extensive and can be difficult to navigate, each participating sector having contributed its own esoteric properties. The number of properties may appear advantageous because this facilitates expressivity. However, their sheer quantity breeds complexity for adopters. In Schema.org’s *Course* specification, for example, there are properties for *Provider* and for *SourceOrganization*. At first sight, although subtly different, these appear to be very similar, and could be used interchangeably if adopters have not understood their distinguishing characteristics. Using different properties to refer to the same thing will clearly have ramifications for search results further down the line. The Bioschemas Training profiles attempt to remedy this by providing marginality categories and reducing the overall options available, confining adopters to unique, clearly distinct terms. The specifications thus offer a more digestible view of Schema.org’s numerous broad vocabularies, filtering out terms that are likely to be irrelevant to the life science training community, and prioritizing those that are likely to be crucial. Hence, for the example mentioned above, the Bioschemas *Course* profile only offers the term *Provider* and not *sourceOrganization*.

Importantly, the Bioschemas Training profiles afford flexibility in terms of how providers can connect related profiles, thereby accommodating different data models. For example, the principal aim of an organization may be to disseminate training materials, using these as the basis for workshops in different locations (*e.g*., The Carpentries). In this instance, where a provider’s data model is lesson-centric, the focus is likely to be on describing the training materials, the courses in which they appear being of secondary importance. Conversely, an organization may focus on providing courses and thus their data model would focus on describing the training opportunities, their associated training materials being of secondary interest.

It is the reciprocal or inverse properties inherent in the Bioschemas Training specifications that allow adopters to annotate their data in line with their own data models. Hence, for example, they can describe their *TrainingMaterials* first, and link to their cognate *Courses* using the *isPartOf* property; or they can describe their *Course* first, and link to the accompanying *TrainingMaterials* with the *hasPart* property. The direction of such links can thus be implemented according to which attributes are considered more critical to a providers’ data model.

In turn, the data annotated using the Bioschemas Training specifications can be harvested by aggregation services like TeSS^24^ (the TeSS team, for example, has developed code that takes the JSON-LD snippets from training providers’ Web pages, extracts the property values used by TeSS’s metadata schema, and creates or updates the requisite registry entry using these values). Such registries could facilitate analyses of archived and current training information from different providers to identify trends or gaps in training provision, and hence inform future training strategies (*e.g*., by targeting under-represented topics or geographic regions). Historic data could also be used by training organizers to improve their event scheduling, identifying and obviating potential clashes with other events in order to maximize the availability of attendees.

One implementation difficulty encountered was with online content and resources: were such resources training materials or courses? Until the Training schemas were developed there was no clear definition, but the semantics of *Course, CourseInstance* and *TrainingMaterial* make this distinction explicit: online learning is defined as an open-ended, asynchronous course with embedded training materials.

Full use of the properties within each profile beyond those with a minimum marginality will be important drivers to the successful interpretation of training courses and materials by search engines and data harvesters. For now, it remains to be seen how search engines will interpret Bioschemas metadata. One particular issue is that search engines may have difficulties determining the current status of training events unless the *eventStatus* property in *CourseInstance* is used. The *eventStatus* property can be used by content providers to indicate that a particular *CourseInstance* has been modified or canceled. When the *eventStatus* property is not used, search engines will have no data with which to render information about the *CourseInstance* correctly. This could lead to a scenario in which, for example, canceled courses still appear to be available.

In the absence of standards like the Bioschemas Training profiles, a lot of work is required to explicitly express the data embedded in websites such that search engines, or other data harvesters, can capture them. Often, content providers will write APIs to express their data. But APIs require substantial work: the developers must design the access points, implement new routes for them, design the resulting data, and write documentation. Data harvesters then have to write custom API clients to interface with the providers’ APIs, essentially repeating work and creating further code-maintenance overheads. The Schema.org implementation upon which Bioschemas is based is simpler than implementing an API endpoint. Moreover, the metadata are encapsulated within websites’ existing pages, so no routing of further access points is required.

Schema.org has an extensive ecosystem of support tools (https://bioschemas.org/developer/software) for creating and validating its annotations. Production-ready validators are being developed for Bioschemas both to make the implementation easier and to verify that the output complies with the extra restrictions involved in the standard, such as marginality, cardinality, CVs, *etc*. Resources for supporting implementations of Bioschemas are thus still needed to ensure that the barriers to adoption remain low.

## Conclusion

The Bioschemas Training profiles initiative set out to make the exchange of training resources easier, breaking down barriers by allowing content providers to more formally structure information on their websites, and effectively ‘speak the same language’. The initiative has led to the articulation of customized Training profiles for describing training information in a standardized way, built on the ubiquitous Schema.org specifications. These profiles have been tailored to the training needs of life scientists following the Bioschemas model, in a manner that helps adopters implement the relevant Training profiles for their content. The hope is that by reducing cognitive and technical overheads, more adopters will be able to implement the standards, thereby facilitating the discovery and exchange of training information.

Although the Bioschemas Training profiles are rooted in the needs of life scientists, the ‘bio’ components of the training specifications are in fact relatively light. There is therefore significant potential for re-use of the specifications, with minimal customisation effort, to represent data across different disciplines.

With four adopters of Course and CourseInstance and eight adopters of TrainingMaterials, the specifications have recently moved from draft to release. We therefore call on the wider training community to embrace and implement these standards as a way to facilitate the global exchange of training information, to make training opportunities and resources easier to discover, and to empower trainers and trainees alike. Their use also embodies the FAIR principles by making training materials and resources Findable-Accessible-Interoperable-Reuseable^25^. Ultimately, we look forward to seeing what new and innovative uses unfold once a web of previously disconnected training data is sewn together to create a more readily navigable and interoperable training landscape.

## Acknowledgements

The authors wish to acknowledge the contributions of members of the Bioschemas Training Profiles Group Members, ELIXIR FAIR Training Focus Group, and The GOBLET Foundation.

They also wish to acknowledge advice received by Phil Barker at LRMI, and the substantial efforts made by the early adopters of these Training profiles, including Bioinformatics.ca, Department of Bioinformatics at Maastricht University, German National Library of Medicine (ZB MED), TeSS ELIXIR-Europe, Bioschemas.org, Galaxy Training, NanoCommons project, EMBL-EBI Training and SIB Swiss Institute of Bioinformatics.

