## Supplementary Figure 1 for "Bioschemas Training Profiles: A set of specifications for standardizing training information to facilitate the discovery of training programs and resources"

### Supplementary Data

**Supplementary Figure 1:** Comparison of training courses with and without Bioschemas Training Profiles standards implementation as validated by Schema.org validator (https://validator.schema.org/). (A) shows the absence of metadata standards to describe the content within a web posted training course in Metabolomics. (B and C) show the use of metadata standards to describe the content within web postings for training courses in Metabolomics. Bioschemas metadata standards markup improves the machine readability of web training resource content.

**(A) CSHL Metabolomics Course 2023**

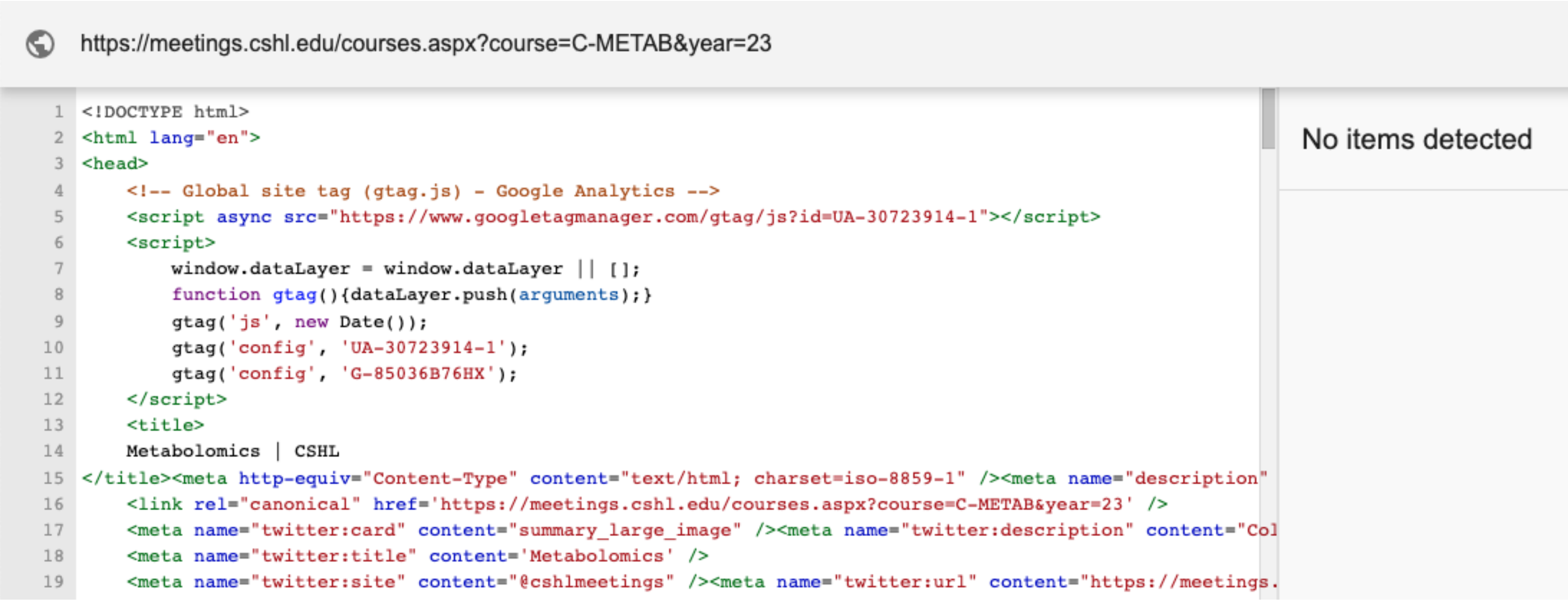

#### (B) Bioinformatics.ca Metabolomics Analysis Workshop 2021

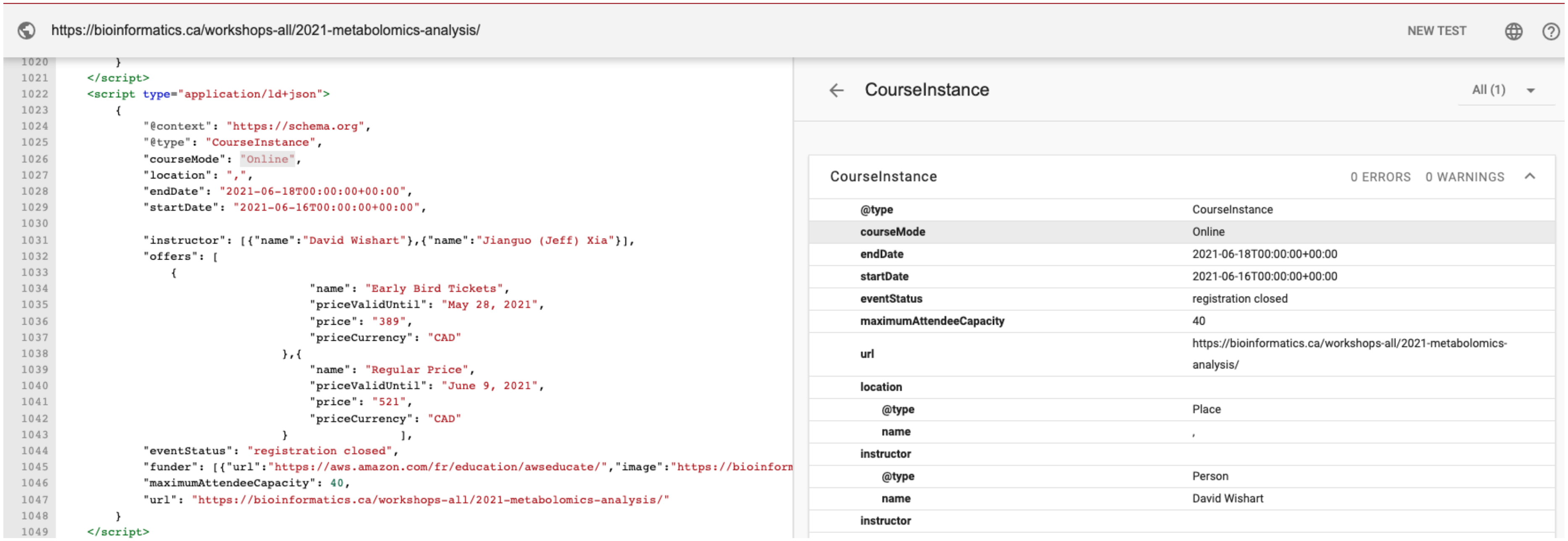

**(C) EBI Introduction to Metabolomics Analysis 2023**

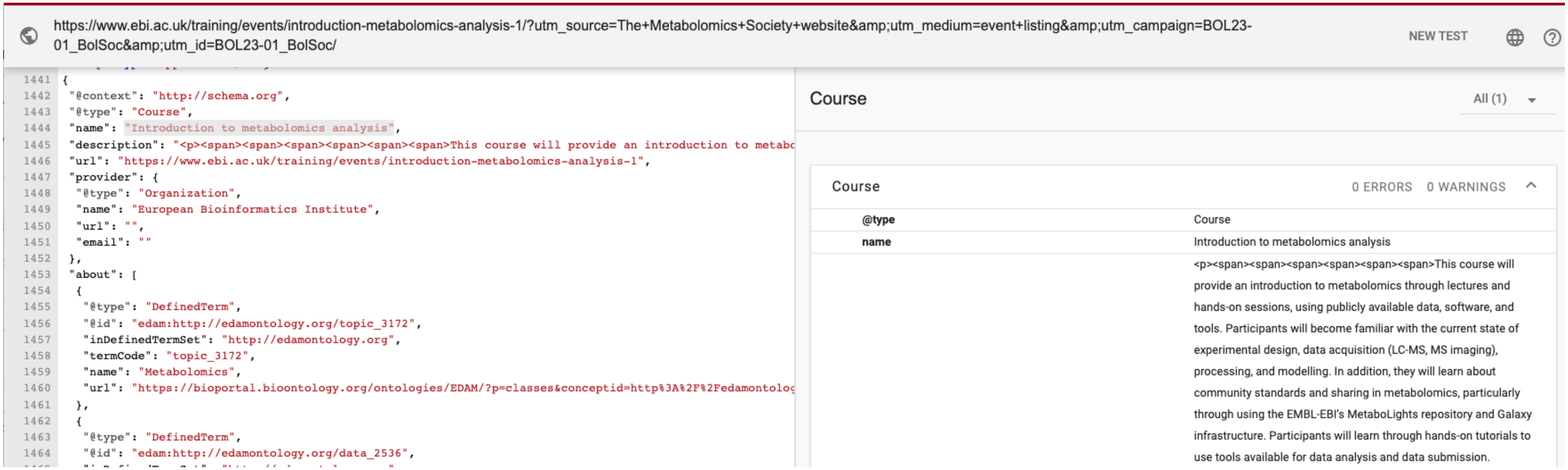
